# *In vitro* disintegration of lozenge in oral cavity

**DOI:** 10.1101/2024.12.17.628961

**Authors:** Quoc Dat Pham, Martin Edman, Katarina Lindell, Marie Wahlgren

**Affiliations:** Department of Food Technology, Lund University, P.O. Box 124, 22100 Lund, Sweden; R&D Product Design, McNeil AB, Box 941, 25109, Helsingborg, Sweden

**Keywords:** release, dissolution, saliva, suction, abrasion

## Abstract

The aim of this study is to develop a new *in vitro* disintegration method for lozenges in the oral cavity (OC) by mimicking physiologically relevant conditions in the OC including saliva flow rate and suction. Disintegration of solid-state dosage forms in the OC can be employed for local absorption in the OC or for delivering to the succeeding parts of the gastrointestinal tract. Understanding the disintegration is important before gaining insight into dissolution and absorption. Here abrasion force and rate exerted on the lozenge during the disintegration are controlled by a texture analyser. We showed that a higher flow rate and more frequent abrasion can lead to a faster disintegration, while variation of the compression force within a range in this study does not affect the disintegration statistically. The results are relevant to different types of consumers with variation in saliva production, for example, people with salivary dysfunction including elderly people. It is also important to consider these effects when formulating flavor and taste of lozenge that can affect consumer perception and consequently suction force and rate or saliva production. We could also observe disintegrated but not dissolved parts of the lozenge by using this apparatus, demonstrating the importance of the correlation between disintegration and dissolution when considering drug absorption in the OC to overcome the challenges of drug clearance from the OC by swallowing.

## 1. Introduction

Oral route is the most frequently used route for drug administration intended for drug absorption through the various epithelia and mucosa of the gastrointestinal tract (GIT). Among different parts along this tract, oral cavity (OC) has been increasingly employed as a release site for solid-state dosage form aiming for local absorption in the OC or for delivering to the succeeding parts of the GIT (Fig. 1A) (Aulton and Taylor, 2018; Rathbone et al., 1994). In addition to the targeted delivery to the OC, the local absorption in the OC offers some advantages compared to the other sites in the GIT, for example, avoiding gastric acid-or digestive enzyme-mediated degradation as well as first-pass effect. The release of solid-state dosage form in the OC (e.g., fast-dissolving or mouth-dissolving tablets and film delivery systems) can also improve convenience of the dosage form e.g., without the need of water and provide a controlled release e.g., throat lozenges. The release profile of solid-state dosage forms in the OC is important to understand the absorption of drug at the desired sites in the GIT and the drug bioavailability. Therefore, there is an increasing demand on *in vitro* – *in vivo* correlations (IVIVC) of the drug release from solid dosage forms to guide early development of a new product and reduce the use of animals and the cost of clinical studies to access bioavailability. The first step for this achievement would be ensuring an IVIVC of the disintegration of the solid dosage form before being able to establish an IVIVC of the dissolution (Fig. 1B).

**Fig. 1.**
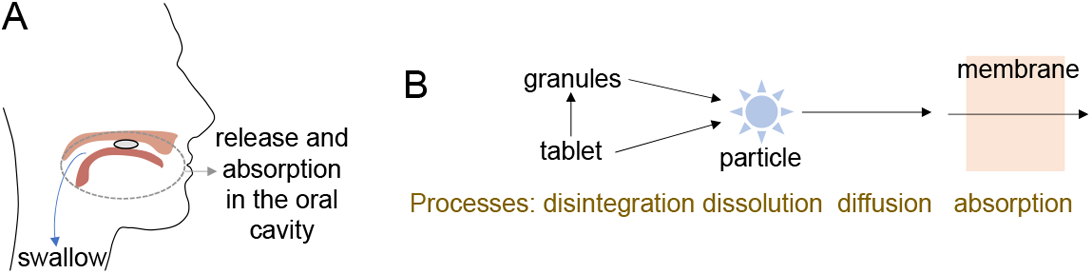
(A) Illustration of lozenge release and absorption in the oral cavity and clearance of drugs from the oral cavity due to swallowing. (B) Different processes during the release and absorption of lozenge in the oral cavity.

IVIVC of the disintegration of tablets in OC has been shown to be not well-correlated in many cases (Abdelbary et al., 2005; Harada et al., 2006; Narazaki et al., 2004) which is likely due to current experimental set-ups which are mainly aimed for tablets dissolved in the stomach. A current disintegration apparatus described in all major pharmacopoeias is a basket-rack assembly that can move vertically, but it does not normally seek to establish an IVIVC (Aulton and Taylor, 2018; Kraemer et al., 2012). Other dissolution testing using stirred-vessel and continuous-flow methods may also provide information on the disintegration time (Aulton and Taylor, 2018). It is noted that the medium volume in these apparatuses is much higher than the accumulated volume of saliva between the swallows (ca 1 mL) (Lagerlof and Dawes, 1984). The volume and the swallow may be critical to the dissolution of chemicals with low solubility and consequently disintegration. Another important factor that is lacking in these apparatuses is abrasion of the tablet surface by hard palate and tongue during sucking process as well as vacuum during swallowing. These actions may be more efficient to erode the tablet surface than the agitation of the medium employed in the compendial dissolution methods, especially for tablets containing gel-forming polymers.

There have been attempts to develop disintegration tests employing mechanical stress force which is static (Abdelbary et al., 2005; Catellani et al., 1989; Dor and Fix, 2000) or dynamic (Harada et al., 2006; Narazaki et al., 2004; Scheuerle et al., 2017; Tietz et al., 2018a; Tietz et al., 2018b), as reviewed in (Al-Gousous and Langguth, 2015; Kraemer et al., 2012). The static mode may not reflect the situation in the OC where the tablet is under peristaltic tongue motion. In some cases when dynamic motion is applied on the tablet, the abrasion force is not controlled and can therefore change simultaneously with the erosion of the tablet. In addition, the tablet may not be entirely in contact with water or the volume and flow of the medium are not physiologically relevant. Learning from the previous studies, we here design and construct a new apparatus (Fig. 2A) that takes into account all the aforementioned aspects to better mimic the conditions in the OC. The new apparatus consists of a flow-through cell in which tablets are hold between a flat solid surface and a plastic sheet. The flow-through cell with a more physiologically relevant medium volume and flow would better mimic the saliva flow in the OC. The cell is mounted on a translational stage and compressed by a force using a texture analyser, imposing controlled pressures and abrasion rates on the tablet to stimulate motion and pressured exerted on the lozenge in the mouth. The controlling of the exerted force during the disintegration is an improvement compared to previous methods (Harada et al., 2006; Narazaki et al., 2004; Scheuerle et al., 2017; Tietz et al., 2018a; Tietz et al., 2018b). The apparatus was employed to test the disintegration of Nicorette^®^ cooldrops 2 mg lozenge (Fig. 2B) that had been studied *in vivo* before so that we can compare the *in vitro* – *in vivo* data. Using this apparatus, we can also learn how different parameters including saliva flow rate as well as abrasion force and rate affect the disintegration, which are relevant to people with salivary dysfunction or dysphagia including elders. These parameters can also vary with lozenge’s flavor and taste that can affect consumer perception. These insights can be utilized to formulate and manufacture lozenges to achieve a better absorption in the OC of different types of consumers.

**Fig. 2.**
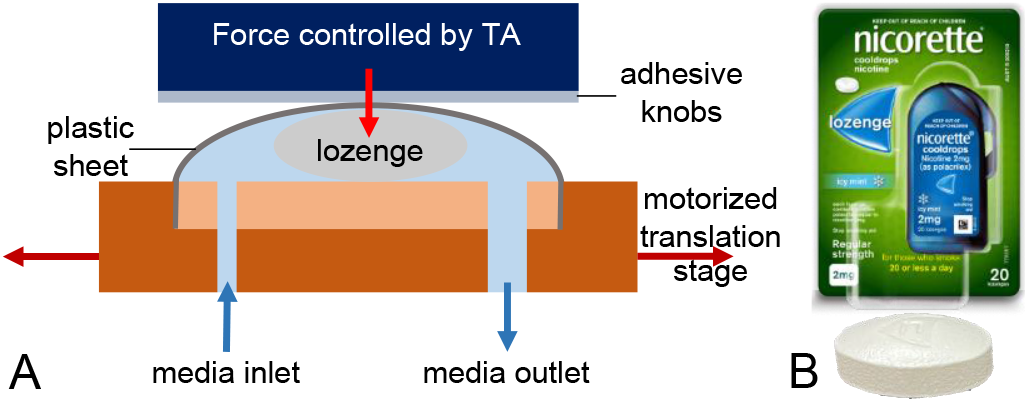
(A) Illustration of the new apparatus for lozenge disintegration in the oral cavity. (B) Nicorette^®^ cooldrops 2 mg tested in this study.

## 2. Materials and methods

### 2.1 Materials

Na2HPO4, KH2PO4 and NaH2PO4 were purchased from Sigma-Aldrich. NaCl was obtained from VWR. Simulated saliva fluid (SSF) pH 6.8 consisting of phosphate buffer saline solution (0.0168 M Na2HPO4, 0.0014 M KH2PO4, 0.0176 M NaH2PO4, and 0.1369 M NaCl) prepared using Milli-Q water with resistivity of 18.2 MΩ cm at 25 °C was used as the medium. The composition is similar to SSF in (Mashru et al., 2005) but NaH2PO4 is added herein to achieve a pH of 6.8 without the need to use phosphoric acid to adjust the pH. The lozenge used is Nicorette^®^ cooldrops 2 mg (Nicotine 2 mg, batch BA165, McNeil Denmark ApS) that has oblong tablet shape with size 14.2 × 9.2 × 6.3 mm (length x width x height) (Fig. 2B).

### 2.2 *In vitro* compendial disintegration experiments

*In vitro* compendial disintegration experiments were examined using DisiTest 50 Disintegration Tester (Dr. Schleuniger^®^ Pharmatron, Switzerland) equipped with a six-tube basket-rack assembly which is immersed in a 1 000 mL beaker containing 800 mL SSF buffer at 37 °C. The size of the glass tubes is 21 mm in diameter and 77 mm in length. One lozenge was placed in each of the six tubes and hold between a wire mesh of the basket-rack assembly and a disc that can detect endpoint of the disintegration automatically. The disintegration of six different lozenges was performed at 30 strokes/min and the length of the stroke is 55 mm.

### 2.3 *In vitro* disintegration using our new apparatus

A flow-through cell was employed in order to mimic the swallow (Fig. 2A). The cell with a base diameter of 20 mm is covered with a plastic sheet (Grippie, PE-LD, b.n.t Scandinavia ab., Sweden) with an area corresponding to a circle with a dimeter of ca 30 mm. The inflow diameter is 1 mm while the outflow is 2 mm which is the same as size of sieves in pharmacopoeial disintegration apparatuses (World Health Organization, 2019). The SSF medium was kept at 37 °C by a hotplate and its flow through the cell is controlled by a Perislastic Pump P-1 (Pharmacia Fine Chemicals, Sweden). The cell is attached to a motorized translation stage 8MT167S-25BS1-Men1, equipped with a stepper and DC motor Controller 1-axis 8SMC5-USB-B8-1 (Standa, Lithuania). The stage was set up to move following this cycle: 1 cm of travel in one direction within ca 1 s and rest for a certain time, and then travel back to the starting position with the same speed and rest for the same period. The constant force applied on the tested lozenge is controlled by a texture analyzer TA-XT2i (Stable Micro Systems, United Kingdom) using a 25 mm diameter cylinder aluminum probe. The force is static during the resting period but can fluctuate around the set value when the stage is moving. Adhesive knobs (b.n.t Scandinavia ab., Sweden) were attached to the probe to hold the plastic sheet while the cell is moving together with the stage. Changes in distance between the probe and the base of the cell were recorded over time, and the disintegration time is defined as when there is no further significant change in the distance (Fig. 3). Experiments with the same set up were performed on 5-6 lozenges, and statistical differences between the disintegration times based on the different experimental set up were determined using two-sample *t*-tests assuming unequal variances with a significance level of 0.05 (Microsoft Excel, Microsoft, United States). A *p* value of < 0.05 was considered to indicate statistical significance.

**Fig. 3.**
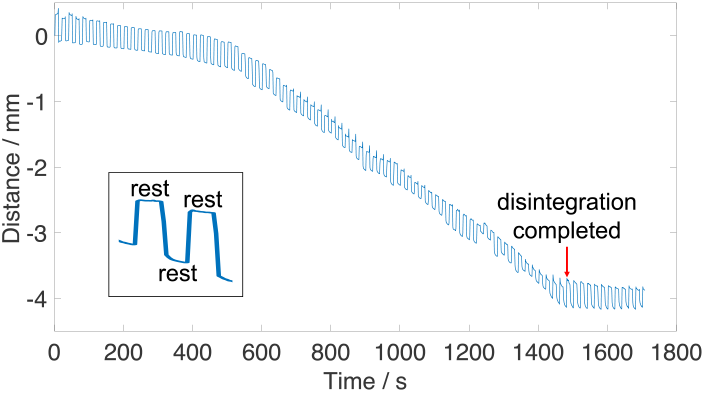
A representative curve of the distance between the probe and the base of the cell versus time recorded at a flow rate of 4 mL/min, a force of 100g and a rest time of 10s. The inset shows the magnification of a part of the curve.

## 3. Results and discussion

In this study the tested lozenge Nicorette^®^ cooldrops 2 mg has *in vivo* intra-oral dissolution time reported to be 16.8 ± 5.5 mins (mean ± SD) (n = 94) ranging from 7 to 37 mins (min-max values) in a clinical pharmacology review, application number NDA 21-330/S-21 (Table 1) (GlaxoSmithKline Consumer Healthcare, 2018). *In vitro* disintegration time of this lozenge recorded using the compendial disintegration tester at a stroke rate of 30 strokes/min is 10.7 ±0.8 mins (mean ± SD) (n = 6) ranging from 9.7 to 12.1 mins (min-max values) (Table 1). It is clear that the mean *in vitro* compendial disintegration time is faster than the mean *in vivo* value. We then performed *in vitro* disintegration tests for this lozenge using our apparatus at different flow rates (2 and 4 mL/min), compression forces (corresponding to 30 and 100 g) and resting periods (10 and 30 s) to mimic multiple potential variations of saliva production rates and suction force and frequency in *in vivo* conditions, respectively. A typical experimental curve of the distance between the probe and the base of the cell is shown in Fig. 3. The pattern of the curve following the set-up cycle of the stage is the result of differences in thickness at the different positions of the lozenge-containing cell (inset in Fig. 3). The probe moved down a distance of ca 4.1 mm when the disintegration finished i.e., no further significant change in the distance. This distance is noted to be smaller than the height of the lozenge of 6.3 mm. This difference is due to the presence of a certain amount of the medium in the cell when the disintegration finishes, while in the beginning the plastic sheet is rather stretched due to the lozenge (Fig. 2A).

**Table 1.** *In vivo* and *in vitro* disintegration time.

|  |  |  |  | Disintegration time (min) |  |  |
| --- | --- | --- | --- | --- | --- | --- |
| | | | | Mean ± SD (n) | Min | Max |
| *In vivo* | | | | 16.8 ± 5.5 (94) | 7 | 37 |
| *In vitro* using the compendial disintegration tester | | | | 10.7 ± 0.8 (6) | 9.7 | 12.1 |
| *In vitro* using the new disintegration apparatus |  |  |  |  |  |  |
| Flow rate (mL/min) | Force (g) | Rest time (s) |  |  |  |  |
| 2 | 100 | 10 | | 29.2 ± 1.4 (5) | 28.0 | 31.5 |
| | 100 | 30 | | 33.7 ± 1.3 (5) | 31.8 | 35.0 |
| | 30 | 10 | | 27.7 ± 0.8 (6) | 27.0 | 29.1 |
| 4 | 100 | 10 | | 25.1 ± 0.8 (6) | 24.3 | 26.6 |
| | 100 | 30 | | 30.2 ± 1.1 (6) | 28.4 | 31.4 |
| | 30 | 10 | | 24.8 ± 1.3 (6) | 22.7 | 26.4 |

We first investigate the effects of flow rate on the disintegration. The stimulated saliva flow rate has been shown to vary quite a lot between individuals, with a range of 2.5 to 4.5 g/min reported for some lozenges (Tietz et al., 2018a) or up to 6.7 mL/min for chewing gums (Dawes and Macpherson, 1992). Here we examine the disintegration at two different flow rates of 2 and 4 mL/min. The results obtained at different experimental settings while keeping the same compression forces (30 or 100 g) and resting periods (10 or 30 s) all showed that a higher flow rate will lead to a statistical faster disintegration (Fig. 4 and Table 1). This is relevant to people with salivary dysfunction, including decreased production of salivary fluid (hyposalivation) and overproduction of saliva (sialorrhea or hypersalivation). Previous studies also showed that saliva production may decrease with aging (Affoo et al., 2015). It is also important to consider the effect of flow rate when formulating lozenge using flavor and taste that can stimulate a significant higher amount of saliva (Watanabe and Dawes, 1988).

**Fig. 4.**
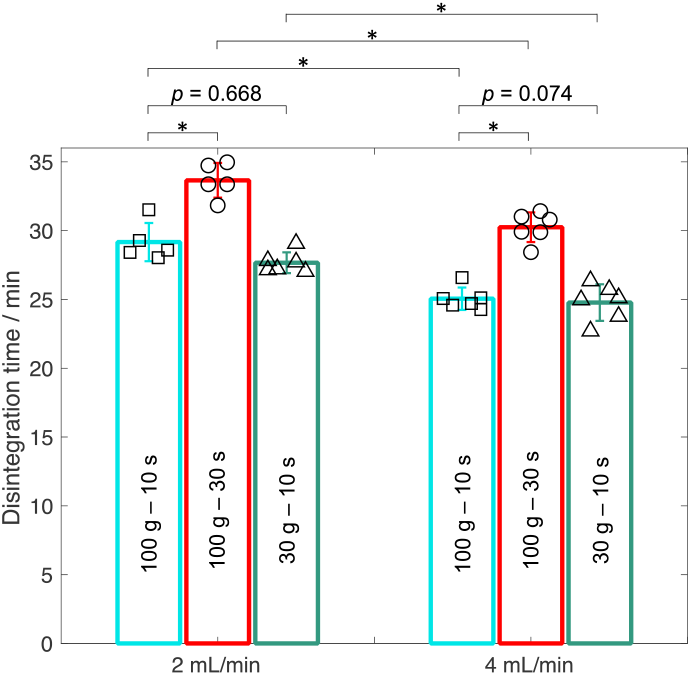
Disintegration time measured at different flow rates (2 and 4 mL/min), compression forces (30 and 100 g) and rest time (10 and 30 s). Data are presented as means ± SD. Significant differences were assessed by two-sample *t*-tests using a significance level of 0.05 and are indicated by * (*p* < 0.0017).

In a next step, we study the impact of the resting period which can be inversely related to the suction frequency. The unstimulated swallowing interval was reported to be around 30-60 seconds (Lagerlof and Dawes, 1984; Rudney et al., 1995) and it is expected to be faster under stimulated conditions. The resting period during the suction is therefore also expected to be shorter than the swallowing intervals since consumer tends to suck at least once before each swallow. Here we choose to study the resting periods of 10 and 30 s, corresponding to 6 and 2 suctions every minute. The disintegration times measured at the same force of 100 g and the same flow rate of 2 or 4 mL/min indicated that a more frequent suction would help the disintegration faster (Fig. 4 and Table 1). The suction frequency would vary between consumer’s suction habit and perception of the lozenge. One can therefore also expect difference in disintegration time for lozenges that differ only in flavor or taste.

Finally, we investigate if the disintegration time is dependent on the compression force exerted on the lozenge during the abrasion. Bolus volume was suggested not to significantly influence maximum lingual force amplitudes during oral stage of swallowing (Miller and Watkin, 1996; Pouderoux and Kahrilas, 1995; Shaker et al., 1988). The force distribution profiles during that swallowing include an increase to maximum force with mean values varying from 63 to 208 g depending on bolus viscosity and then a decrease in force until a sudden return to resting state is obtained (Miller and Watkin, 1996). In addition, lingual swallowing pressure was shown to vary significantly with taste of bolus (Pelletier and Dhanaraj, 2006) or being lower in people with physical impairment to swallowing (dysphagia). Abdelbary et al. (Abdelbary et al., 2005) studied *in vitro* disintegration of rapidly disintegrating tablets using a texture analyser apparatus which is set to maintain a predetermined nominal force of 50 g. Here we use a compression force of 30 g or 100 g during the whole experiments to mimic multiple potential average suction forces. The results in Fig. 4 and Table 1 show that there is no significant difference (*p* > 0.05) in the disintegration time resulted from increasing the compression force from 30 to 100 g while keeping the same flow rate (2 or 4 mL/min) and resting period of 10 s. In another *in vitro* study on oral rapidly disintegrating tablets in the mouth during breastfeeding, compression level was found to be a significant factor for increasing the release at rotational rates representing nonnutritive breast-feeding (Scheuerle et al., 2017). These differences can be due to the types of tablets and compression forces during tableting which is linked to the fact that the oral rapidly disintegrating tablets are expected to be more sensitive to the abrasion force since they should disintegrate much faster.

The *in vitro* disintegration time recorded at the different experimental settings using the new apparatus varies between 22.7 and 35.0 mins which is in the range of the *in vivo* intra-oral dissolution time from 7 to 37 mins (min-max values) (Table 1) (GlaxoSmithKline Consumer Healthcare, 2018). Still, the mean *in vivo* value of 16.8 ± 5.5 mins is lower than the mean *in vitro* values in Table 1. As shown in the previous paragraphs in this study, a higher flow rate and/or a more frequent abrasion (as used in previous studies) (Tietz et al., 2018a; Tietz et al., 2018b) can lead to a faster disintegration, matching the *in vivo* value. It may not be meaningful to capture the mean *in vivo* disintegration time of different individuals that have a big variation in saliva production rate and suction force and rate by an *in vitro* experimental setting of flow rate, compression force and resting period, and these parameters may not change proportionally with the disintegration time. Here we showed that using the certain experimental settings that are physiologically relevant we could obtain the *in vitro* disintegration time within the range of the *in vivo* data and learn how these parameters affect the disintegration. It is also noted that the *in vitro* compendial disintegration time is much faster than our recorded values with the new apparatus (Table 1) which can be due to the strokes at a faster rate (30 strokes/min) and also the larger volume of the medium (800 mL).

We also noted that during the experiments we could visually observe solid particulates that are pumped out of the cell through the media outlet, clearly indicating that a fraction of the disintegrated lozenge is not dissolved within the cell. One may need an in-situ (in-line) measurement right at the beginning of the media outlet in order to quantify the amount of the undissolved fraction and this is outside the scope of this study. These disintegrated but not dissolved parts would be likely swallowed in *in vivo* condition, therefore reducing the absorption and bioavailability in the OC. One may therefore need to optimize the dissolution rate to ensure all the disintegrated parts are dissolved before being swallowed. This can be done by reformulating the formulation (e.g., choosing smaller particle size) or lowering compression forces during tableting while still maintaining the optimized disintegration. Shape and size of lozenge were also suggested to impact the *in vivo* sucking time (Tietz et al., 2018a). On the other hand, one can slow down the disintegration rate if the concentration of drug in plasma is still within its therapeutic window. There are also cases when one wants to employ lozenges to deliver drugs to stomach or small intestine rather than absorption in the OC, and the disintegration should be maximized while the dissolution should be limited.

## 4. Conclusion

Using the new apparatus that can mimic the physiologically relevant conditions in the OC including flow rate, compression force and resting period, we can obtain the *in vitro* disintegration time of the lozenge within the range of the *in vivo* data. In addition, we showed that a higher flow rate and more frequent abrasion can lead to a faster disintegration, while the variation of the compression force (between 30 and 100 g) does not affect the disintegration statistically. These results would be useful when formulating flavor and taste of lozenge that can affect consumer perception and consequently suction rate or saliva production. One may also expect different disintegration times when the lozenge is consumed by people with salivary dysfunction including elderly people. Finally, the observation of disintegrated but not dissolved parts of the lozenge by using this apparatus illustrates the importance of the correlation between disintegration and dissolution when considering drug absorption in the OC to overcome the challenges of drug clearance from the OC by saliva. Future studies using higher compression force (e.g., when consumers are engaged in the lozenge) or employing other types of lozenges could be performed.

## Acknowledgements

This work was supported by McNeil AB, Helsingborg, Sweden. We kindly acknowledge Per Steen and Hans Bolinsson for help with setting up the experiments.

## Appendix A

**Fig. A1.**
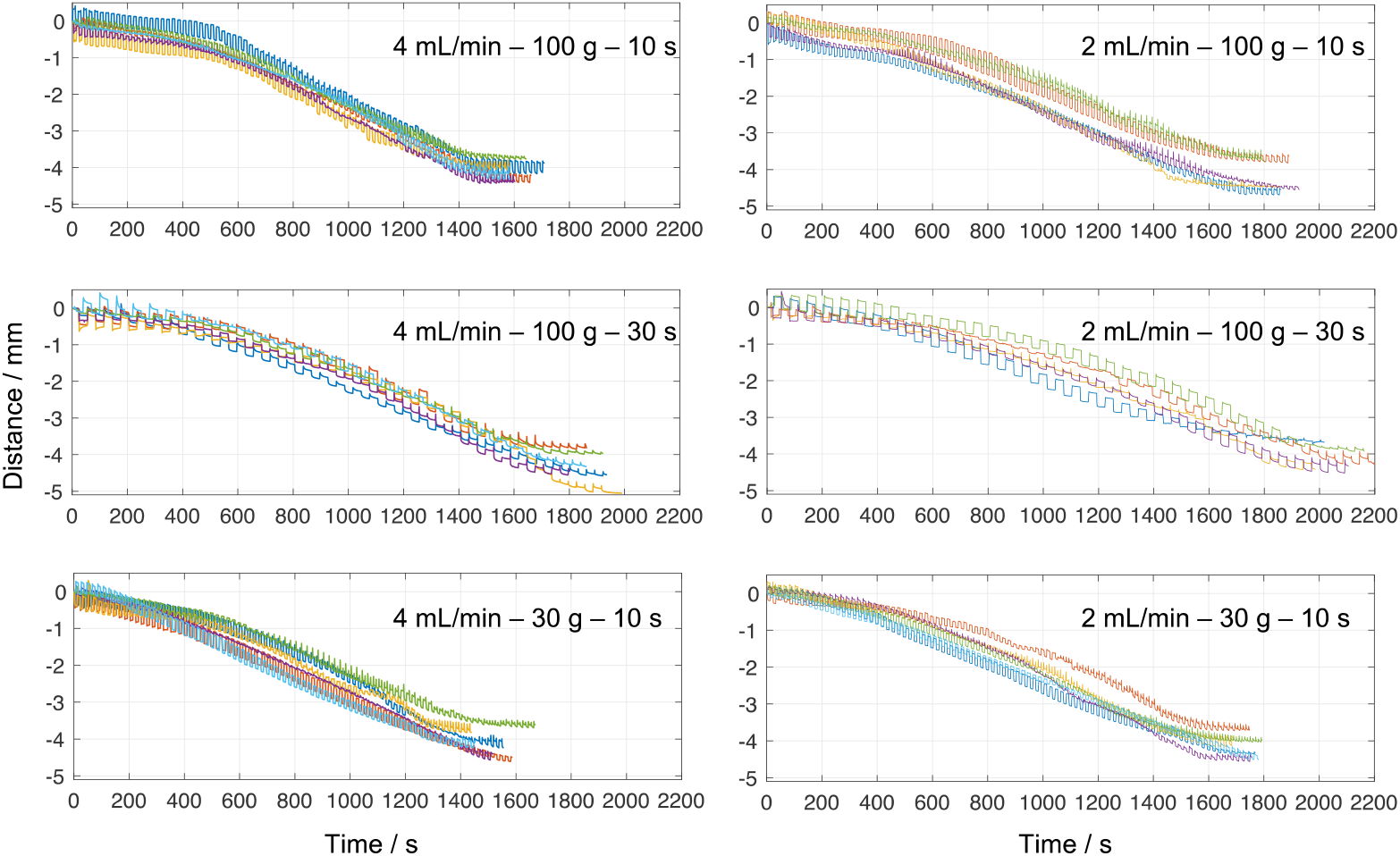
Distance between the probe and the base of the cell versus time recorded at different experimental settings of flow rate, compression force and rest time.

